# Genome-wide analysis reveals pathways important for the development and maturation of excitatory synaptic connections to GABAergic neurons

**DOI:** 10.1101/2025.07.05.663213

**Authors:** Devyn B. Oliver, Shankar Ramachandran, Kasturi Biswas, Claire Y. Bénard, Maria Doitsidou, Hailey McKillop, Noelia Genao, Michele L. Lemons, Michael M. Francis

## Abstract

A high degree of cell and circuit-specific regulation has presented challenges for efforts to precisely define molecular mechanisms controlling synapse formation and maturation. Here, we pursue an unbiased forward genetic approach to identify *C. elegans* genes involved in the formation and maturation of cholinergic synaptic connections with GABAergic motor neurons as indicated by the distribution of GFP-tagged postsynaptic acetylcholine receptors (AChR) on GABAergic dendrites. We identified mutations in 3 genes that identify key processes in synapse/circuit maturation. Mutation of the RUN domain (<u>R</u>PIP8, <u>U</u>NC-14, and <u>N</u>ESCA) cargo adaptor gene *unc-14* dramatically impacts overall GABAergic neuron morphology and dendritic spines. Mutation of the nicotinic acetylcholine alpha subunit gene *unc-63* causes a failure in AChR assembly in GABAergic neurons but does not significantly alter dendritic spine structure or abundance. Finally, a mutation in the Liprin-α synaptic scaffold gene *syd-2* severely disrupts both dendritic spines and AChR localization. The identification of these three genes from our screen highlights how mechanisms for cargo trafficking, receptor assembly, and synapse structural organization each make distinct contributions to synapse assembly and circuit connectivity.

## Introduction

Nervous system performance relies on the precise organization of synaptic connections between neurons. During synapse formation, presynaptic and postsynaptic machinery must be properly trafficked to nascent sites of synaptic contacts and stabilized. Presynaptic protein complexes recruit neurotransmitter-filled synaptic vesicles to specific sites, known as active zones (AZ), and neurotransmitter receptors are clustered at high density in apposition to these AZs. Specific neurotransmitter receptor types are localized in a cell- and synapse-type specific manner, based at least in part on their subunit composition. While we now have generalized knowledge of the various processes that guide these steps in synapse development, the molecular mechanisms involved often show a high degree of cell-type specificity, complicating efforts to gain a comprehensive mechanistic understanding of synapse development processes across neuron classes and neurotransmitter systems.

Genome-wide screening approaches in invertebrates have proven particularly valuable for identifying conserved genetic pathways involved in synapse formation. In *C. elegans*, these efforts have largely focused on the identification of genes involved in the organization of neuromuscular synapses. For example, screens to identify mutants with mislocalized synaptic vesicle markers at GABAergic neuromuscular synapses identified conserved genes such as *syd-2* that encode presynaptic scaffolds important for active zone structural organization [1]. Additionally, screens to identify mutants resistant to the effects of the ionotropic (nicotinic) acetylcholine receptor (AChR) agonist levamisole identified extracellular scaffold genes important for the clustering of muscle AChRs at cholinergic neuromuscular synapses [2,3]. By comparison, molecular mechanisms critical for the formation of synapses onto *C. elegans* neurons are less well defined. Our previous work showed that the presynaptic adhesion protein NRX-1 stabilizes developing postsynaptic spines on GABAergic dendrites [4,5]. However, additional players in the development of spines and their associated postsynaptic sites have remained unidentified.

In this study, we pursued a genome-wide screening approach to obtain mutants in which the organization of postsynaptic cholinergic receptor clusters (AChRs) on GABAergic motor neurons is disrupted. From this screen, we obtained three mutants that indicate molecular mechanisms important for the development and maturation of excitatory synaptic contacts located on dendritic spines of GABAergic motor neurons. We isolated a new mutant allele of the *unc-14* gene that disrupts both synaptic organization and dendrite outgrowth. *unc-14* encodes a RUN domain adaptor protein previously implicated in axonal trafficking and cargo selection [6-9]. Further, we isolated a new mutant allele of the nicotinic receptor alpha subunit *unc-63* that disrupts localization of ACR-12 receptor clusters in GABAergic dendrites, providing evidence that UNC-63 is an obligate component of ACR-12-containing receptors. Finally, we isolated a new allele of the synaptic scaffold *syd-2*/Liprin-α, previously shown to be important for the organization of presynaptic sites [1,10,11], that disrupts the outgrowth of dendritic spines and the clustering of postsynaptic receptors in GABAergic neurons. Our evidence linking these genes with synapse formation in *C. elegans* GABAergic neurons advances a new model for analysis of their functional contributions.

## Results

### A forward screen to identify genes important for neuronal AChR clustering/assembly

To identify genes important for the formation or maintenance of synaptic connections in the *C. elegans* motor circuit, we performed a forward genetic screen to obtain mutants where the clustering and/or distribution of AChRs in dorsal type D class (DD) GABAergic motor neurons is disrupted. ACR-12-containing AChR clusters are organized into synapses located at the tips of spines in mature DD GABAergic dendrites [4,5,12,13]. We screened L4 stage F2 progeny of EMS-mutagenized animals stably expressing ACR-12::GFP in GABAergic neurons [14,15] for mutants with either misplaced or altered density of AChR clusters in the dendrites of DD GABAergic neurons. We identified three mutant alleles of note: *uf169*, *uf175*, and *uf174*. For each of these mutant alleles, we noted striking decreases in ACR-12::GFP fluorescence in GABAergic dendrites. Using next generation sequencing and variant discovery mapping, we identified the causal mutations in these 3 candidates and describe our analysis of each below.

### A mutation in the *unc-14*/RUN domain cargo adaptor disrupts GABAergic neurite outgrowth and organization

Whole genome sequence analysis and variant discovery mapping of *uf169* mutants revealed a point mutation that introduces an early stop in the first exon of the *unc-14* gene, presumably resulting in a null allele (**Fig. 1A,B**). UNC-14 encodes a RUN domain (<u>R</u>PIP8, <u>U</u>NC-14, and <u>N</u>ESCA) cargo adaptor protein [6]. RUN domain-containing proteins associate with small GTPases, motor proteins, and cytoskeletal networks, and are implicated in a variety of cellular signaling pathways and processes, including neuronal development [16,17]. UNC-14 was previously implicated in the outgrowth of GABAergic axons through interactions with the UNC-51 serine/threonine kinase and Netrin signaling components [6,7,9,18,19] and synaptic vesicle transport through interactions with kinesin motors UNC-116 and KLC-2 [8], but its importance in the extension of GABAergic dendrites and the structural organization of postsynaptic sites has not been previously investigated.

**Figure 1.**
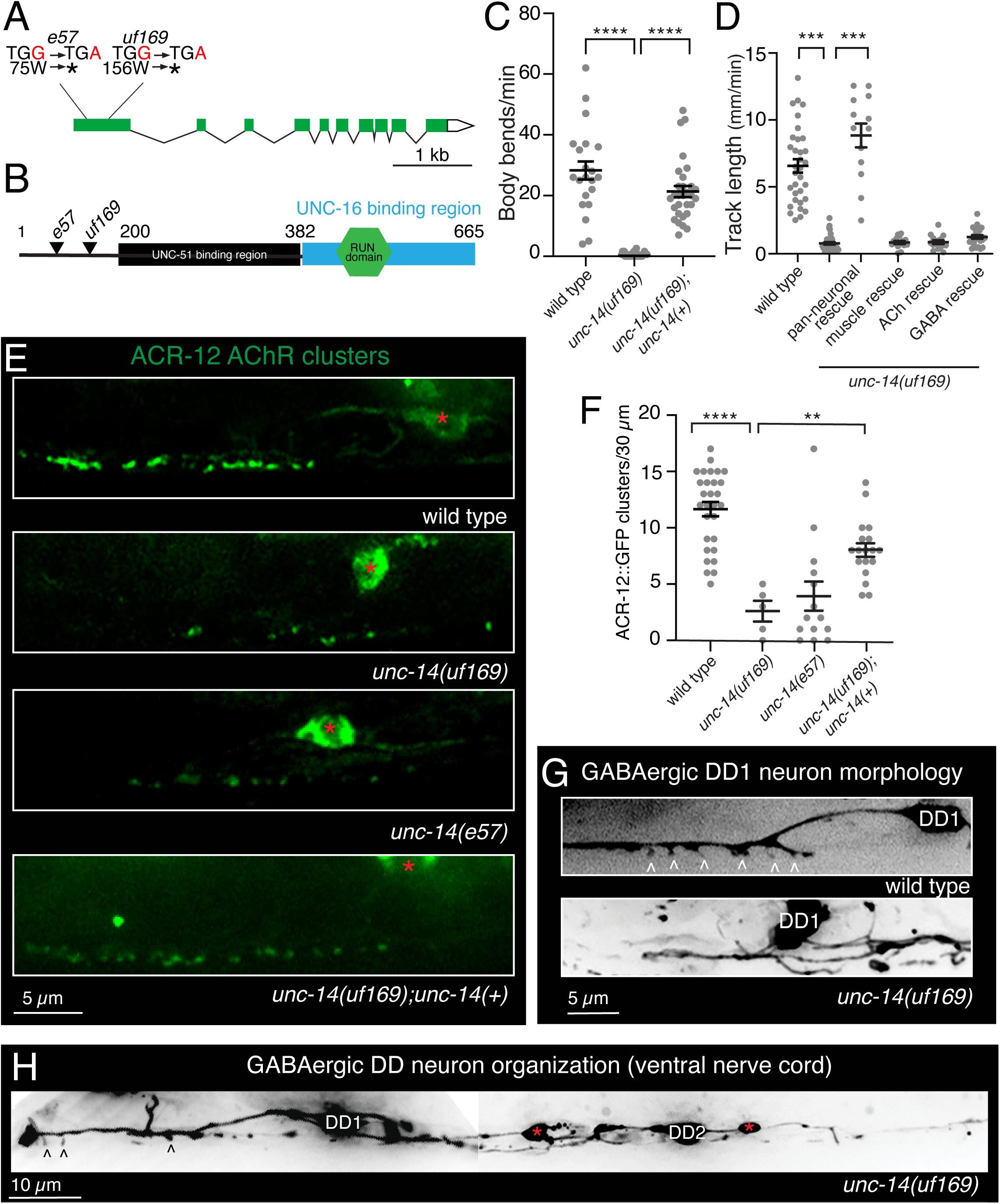
*unc-14* mutation disrupts the outgrowth of GABAergic dendrites. **A.** Schematic of *unc-14* transcript. Reference allele (*e57*) and screen isolate allele (*uf169*) mutations are indicated, both resulting in a premature stop (*) in the first exon. **B.** UNC-14 protein domains including the predicted binding sites of UNC-51 and UNC-16. Modified from [8]. Reference allele (*e57*) and screen isolated allele (*uf169*) mutations are indicated. Numbers, amino acid positions. **C.** Quantification (mean ± SEM) of the average body bends per minute during exploratory movement for the indicated genotypes. Points indicate measurements from independent animals. \*\*\*\**p*<0.0001, one-way ANOVA with Dunnett’s multiple comparison’s test. N=22,32,30. **D.** Quantification (mean ± SEM) of track length (mm/min) during exploratory movement for the indicated genotypes. The decreased movement of *unc-14(uf169)* mutants was rescued by pan-neuronal (P*rgef-1) unc-14(+)* expression but not by cholinergic (P*unc-17*β), GABAergic (P*unc-47*) or muscle (P*myo-3*) *unc-14(+)* expression in *unc-14(uf169)* mutants. Points indicate measurements from independent animals. \*\*\**p*<0.001, one-way ANOVA with Dunnett’s multiple comparison’s test. N=33,42,13,15,21,25. **E.** Representative confocal images of ACR-12 AChR localization (P*flp-13*::ACR-12::GFP) in ventral GABAergic DD1 dendrites of control, screen isolate *unc-14(uf169)*, *unc-14(e57)*, and *unc-14(uf169)* rescued by expression of wild type *unc-14* coding sequence and promoter region. Red asterisks, DD1 cell body. **F.** Quantification (mean ± SEM) of the number of ACR-12 AChR clusters in a 30 µm synaptic region of the DD1 dendrite in the ventral nerve cord of wild type, screen isolate *unc-14(uf169)*, *unc-14(e57)*, and *unc-14(uf169)* rescued by expression of wild type *unc-14* coding sequence and promoter region. Points indicate measurements from independent animals. \*\*\*\**p*<0.0001, \*\**p*<0.01, one-way ANOVA with Dunnett’s multiple comparison’s test. N=29,5,14,18. **G.** Representative confocal images of DD1 dendrite in the ventral nerve cord (P*flp13*::mCherry) of wild type and *unc-14(uf169)* mutants. White carets indicate DD dendritic spines. **H.** Representative image showing expanded view of the ventral nerve cord processes of DD1 and DD2 GABAergic neurons in the *unc-14(uf169)* mutant. Black carets point to possible DD dendritic spines. Red asterisks indicate varicosities along the ventral nerve cord.

*unc-14(uf169)* mutants show severely uncoordinated movement (**Fig. 1C**), similar to previously characterized *unc-14(e57)* loss-of-function mutants. Expression of the wild-type *unc-14* coding sequence and 2.9 kb promoter region upstream of the *unc-14* start in *unc-14(uf169)* mutants robustly restored normal movement (**Fig. 1C**), supporting that *unc-14(uf169)* is a loss-of-function mutation. Pan-neuronal, but not muscle, expression of wild-type *unc-14* was also sufficient to rescue the locomotor deficits of *unc-14(uf169)* mutants (**Fig. 1D**), indicating a neuronal requirement. However, specific expression in either GABAergic or cholinergic neurons was not sufficient for behavioral rescue (**Fig. 1D**), consistent with a broad neuronal requirement for *unc-14* expression.

We next quantified the effects of the *uf169* mutation on ACR-12::GFP fluorescence. We observed the number of fluorescent receptor clusters is severely reduced in DD GABAergic dendrites of *uf169* mutants (**Fig. 1E,F**). Dendritic ACR-12::GFP fluorescence is similarly reduced in the DD dendrites of previously characterized *unc-14(e57)* loss-of-function animals (**Fig. 1E,F**). The reduction in ACR-12/AChR clusters was rescued by expression of the wild-type *unc-14* sequence (2.9 kb promoter) (**Fig. 1E**). Notably, we also observed an increase in the ACR-12::GFP fluorescence in DD neuronal somas of *unc-14(uf169)* mutants relative to the unmutagenized starting strain, perhaps indicative of impaired trafficking from the soma. To better understand how mutation of *unc-14* impacts AChR localization we examined the morphological features of DD neurons in *unc-14* mutants expressing a P*flp-13*::mCherry transcriptional reporter. We observed a significant reduction in dendritic spines in *unc-14(uf169)* mutants (**Fig. 1G**). However, this reduction was accompanied by severe morphological defects in ventral nerve cord DD neurites, including ectopic branching, large varicosities, and misplaced DD somas in the *uf169* mutant (**Fig. 1H**). Given the severe dendrite morphology defects observed in these animals, the reduction in spine number likely arises from a generalized requirement for UNC-14 in dendrite extension rather than a specific role for UNC-14 in dendritic spine outgrowth. While *unc-14* has previously been shown to play a role in neurite outgrowth and extension [6,9,19], our data suggest an additional role for UNC-14 in postsynaptic receptor trafficking and localization.

### A mutation in the nAChR alpha subunit *unc-63* gene disrupts ACR-12 AChR clustering in DD motor neurons

*uf175* mutants displayed a striking reduction in AChR clusters in GABAergic dendrites. Sequence analysis and mapping of *uf175* mutants revealed a missense mutation in exon 2 of the *unc-63* gene (**Fig. 2A**). *unc-63* encodes a nicotinic acetylcholine receptor (nAChR) alpha subunit that is expressed in both the nervous system and muscles, and is incorporated into heteromeric nAChRs that vary in subunit composition across cell types [5,20-24]. The sensitivity of UNC-63-containing nAChRs at the *C. elegans* neuromuscular junction to the nicotinic agonist levamisole (L-AChR) has been well-documented [25,26]. UNC-63 also contributes to other heteromeric nAChR classes in both cholinergic and GABAergic motor neurons [5,20,23]. The missense mutation carried by *unc-63(uf175)* mutants introduces a P44L substitution at a highly conserved position in the N-terminal extracellular region of UNC-63 immediately adjacent to amino acids implicated in forming the main immunogenic region (MIR) of the human muscle alpha1 nAChR subunit (**Fig. 2B**). The MIR is a major target for autoantibodies in humans suffering from myasthenia gravis [27,28]. Sequence in this region is thought to be important for mediating the conformational maturation of AChRs during receptor assembly as well as for determining sensitivity to activation by ACh [29].

**Figure 2.**
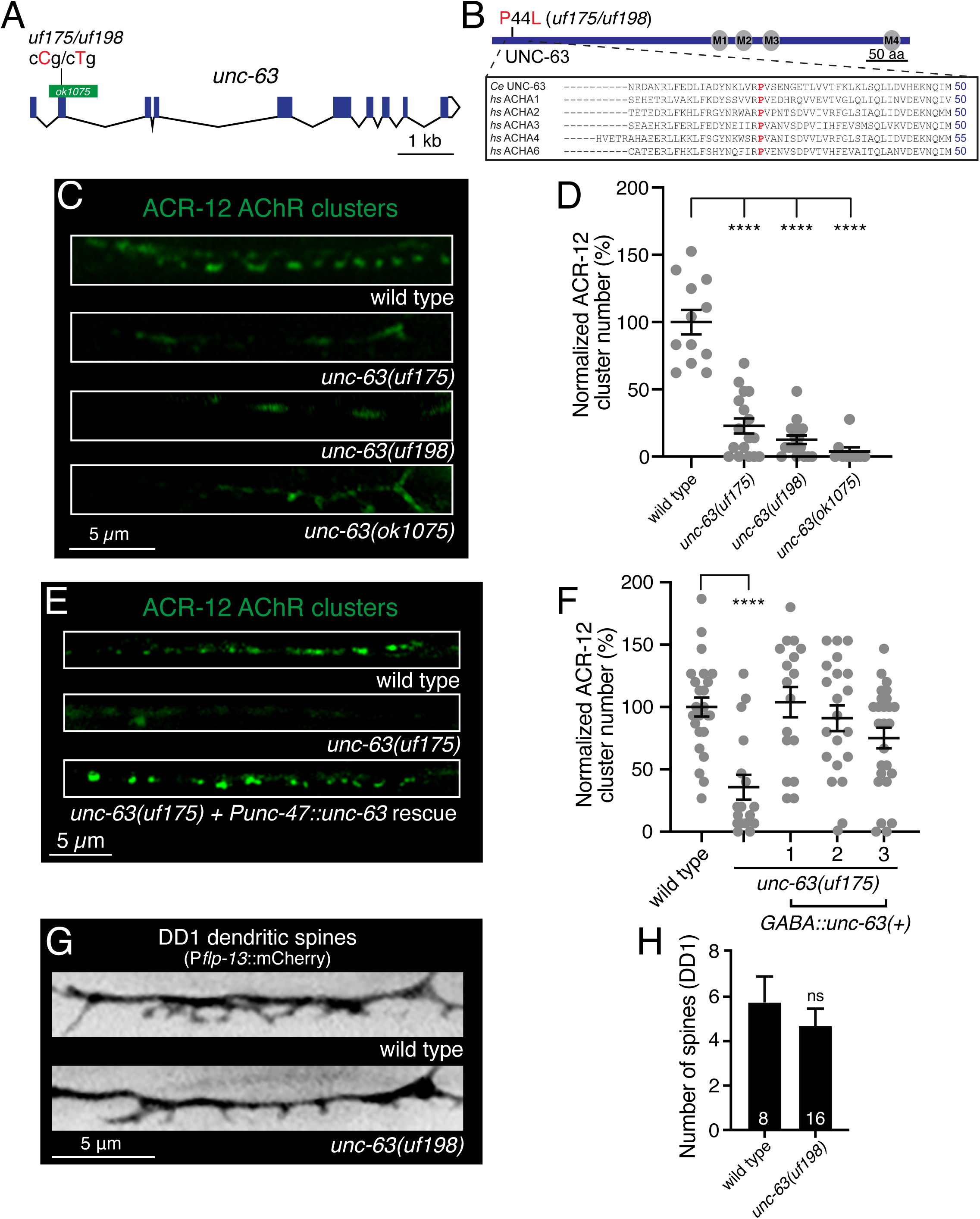
Substitution of a highly conserved proline residue near the N-terminus of the UNC-63 AChR subunit disrupts ACR-12 AChR localization in GABAergic DD dendrites. **A.** Transcript of *unc-63* with annotated *unc-63* alleles. Screen isolated *uf175* and CRISPR generated *uf198* alleles are indicated. Green bar indicates region *unc-63(ok1075)* deletion. **B.** Protein domain structure of UNC-63. Transmembrane domains M1 through M4 are indicated. Screen isolated *uf175* and CRISPR generated *unc-63(uf198)* P44L mutation are shown. Sequence alignments of UNC-63 across species are detailed in box. The red “P” indicates the *unc-63(uf198*) P44L mutation, a proline residue at this position is highly conserved from *C. elegans (Ce)* to humans (*hs*). **C.** Confocal images of ACR-12 localization (P*flp-13*::ACR-12::GFP) in GABAergic DD1 dendrites of wild type, *unc-63(ufI175)*, *unc-63(uf198)* and *unc-63(ok1075)* animals. **D.** ACR-12 clusters in GABAergic dendrites are decreased by mutation of *unc-63*. Quantification of receptor clusters, normalized to wild type, for each genotype. Points indicate measurements from independent animals. Mean ± SEM are indicated. \*\*\*\**p*<0.0001, one-way ANOVA with Dunnett’s multiple comparisons test. N=12,17,17,9. **E.** Confocal images of DD1 ACR-12 localization (P*flp-13*::ACR-12) in) in GABAergic DD1 dendrites of wild type, *unc-63(uf175),* and *unc-63(uf175)* rescued by GABA-specific (P*unc-47::unc-63 cDNA*) expression of wild type *unc-63*. **F.** GABA-specific (P*unc-47::unc-63 cDNA*) expression of a wild type *unc-63* cDNA in *unc-63(uf175)* mutants restores ACR-12 clustering (3 independent lines). Quantification of receptor clusters, normalized to wild type, for each genotype. Points indicate measurements from independent animals. Mean ± SEM are indicated. \*\*\*\**p*<0.0001, one-way ANOVA with Dunnett’s multiple comparisons test. N=24,17,17,21,26. **G.** Representative confocal images of DD1 spines (P*flp-13*::mCherry) in wild type and *unc-63 (uf198)* animals. **H.** Quantification of spine number (mean ± SEM) in DD1 dendrites (P*flp-13*::mCherry) of wild type and *unc-63(uf198)* animals. are indicated. ns, not significant. Numbers in bars indicate the number of animals quantified.

ACR-12 AChR clusters in the DD neurons of *unc-63(uf175)* mutants are decreased by ∼65% compared with wild type (**Fig. 2C,D**). Mutant *unc-63(uf198)* animals, carrying the same cCg/cTg P44L mutation engineered by CRISPR/Cas9 genome editing, showed a similar decrease in ACR-12 AChR clusters (**Fig. 2C,D**), providing additional evidence that the P44L substitution is the causal mutation. Similar decreases in ACR-12 clustering were also observed in previously characterized *unc-63(ok1075)* animals (**Fig. 2C,D**) that carry a deletion in the *unc-63* locus [5], suggesting the *uf175* mutation isolated from the screen and the corresponding CRISPR-generated *uf198* mutation cause a severe loss-of-function. Cell-specific expression of wild-type *unc-63* in the GABAergic neurons of *unc-63(uf175)* mutants rescued the decrease in ACR-12 AChR clusters (**Fig. 2E,F**), demonstrating a cell-autonomous requirement for UNC-63 in AChR assembly and synaptic clustering in DD neurons. In contrast to the severe decrease in AChR clusters, the number of dendritic spines in DD neurons is not appreciably altered by the UNC-63 P44L substitution (**Fig. 2G,H**), consistent with our prior findings that spine formation and stabilization occur independently of postsynaptic receptor localization [4,5].

### The UNC-63 P44L substitution is semi-dominant and disrupts neuromuscular function

In addition to motor neurons, *unc-63* is strongly expressed in body wall muscles and serves an obligate subunit of heteropentameric postsynaptic levamisole-sensitive L-AChRs at neuromuscular synapses [21]. We next investigated the function of muscle L-AChRs in *unc-63(uf198)* mutants to explore how the P44L substitution may impact classes of UNC-63 containing AChRs in other cell types. Our interest in this question was motivated in part by our initial qualitative observation that the movement of *unc-63(uf198)* animals appeared phenotypically less severely affected compared with strains carrying the previously characterized *unc-63(x37)* loss-of-function mutation [22]. To investigate this possibility, we measured movement velocity by video monitoring and post-hoc analysis of worm locomotion (**Fig. 3A**). We found that the *unc-63(x37)* loss-of-function significantly decreased the movement velocity as previously reported [22]. In contrast, the P44L substitution in *unc-63(uf198)* mutants produced no appreciable decrease in movement velocity. To explore this more deeply, we investigated the effects of *unc-63* mutation in combination with deletion of the *acr-16* gene. *acr-16* encodes another nAChR alpha subunit that forms homomeric iAChRs (N-AChR) which operate in parallel with L-AChRs at muscle synapses. Combined loss of function of these two receptor classes produces severe deficits in both cholinergic neuromuscular transmission and movement [30,31]. Consistent with prior work, we found that the movement of *unc-63(x37);acr-16(ok789)* double mutants is severely decreased compared with either wild type or *acr-16(ok789)* single mutants. In contrast, the movement *unc-63(uf198);acr-16(ok789)* double mutants is not significantly decreased (**Fig. 3A**). Together, our results suggest there may be sufficient muscle L-AChR function in *unc-63(uf198)* mutants to support muscle contraction and movement.

**Figure 3.**
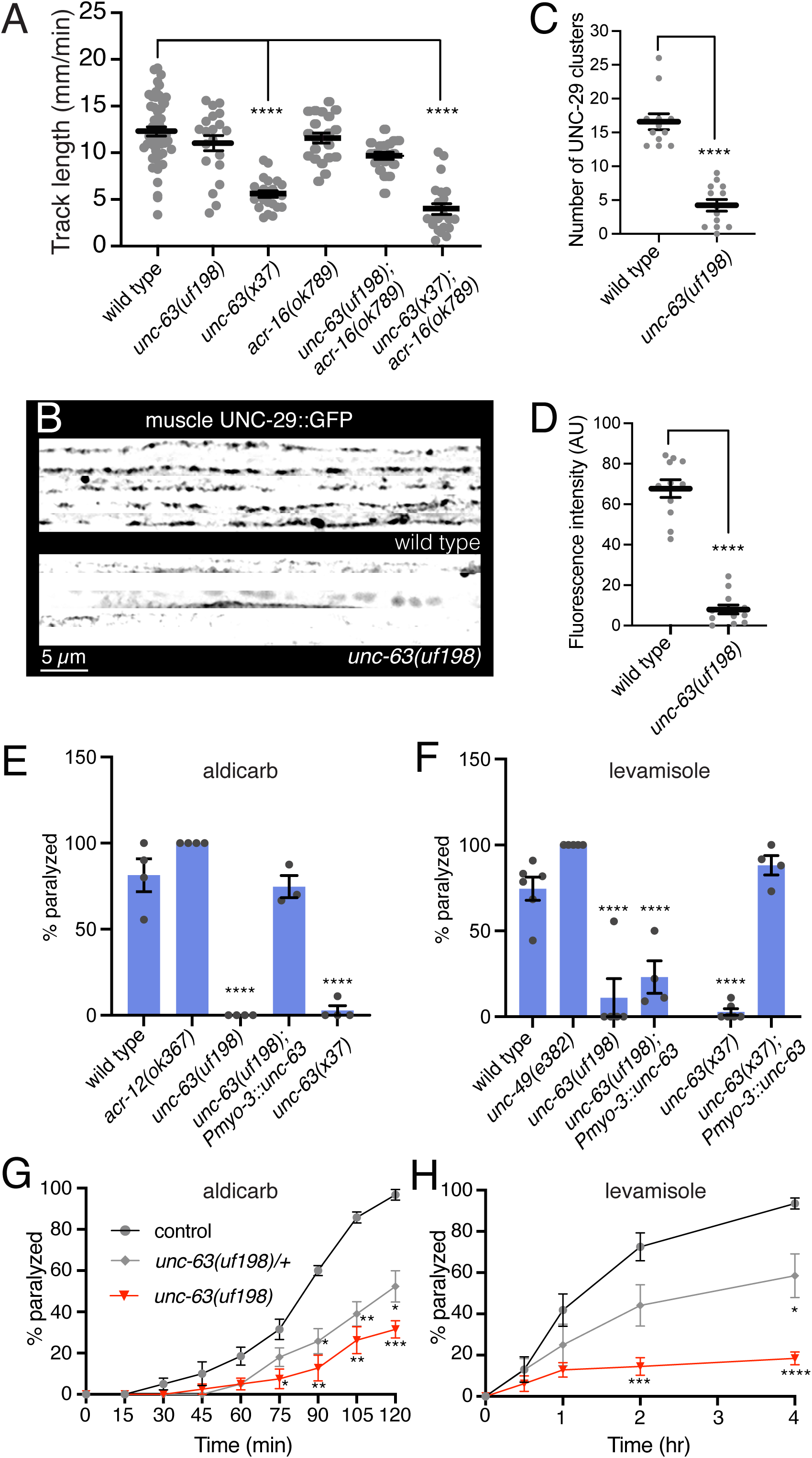
The UNC-63 P44L substitution disrupts neuromuscular synapse organization and presents resistance to aldicarb and levamisole. **A.** Quantification of track length (mm/min) during exploratory movement. *unc-63(uf198)* mutants move similarly to wild type while the movement of *unc-63(x37)* mutants is more severely disrupted. Similar results are obtained in animals that also lack functional ACR-16 N-AChRs. Points indicate measurements from independent animals. Mean ± SEM are indicated. \*\*\*\**p*<0.0001, one-way ANOVA with Dunnett’s multiple comparisons test. N=52,22,20,21,22,19. **B.** Representative confocal images of muscle regions along the ventral nerve cord from five wild type and *unc-63(uf198)* mutants expressing P*myo-3*::UNC-29::GFP (muscle AChRs). **C,D.** UNC-29::GFP AChR clusters (C) and muscle fluorescence intensity (D) are each significantly decreased in *unc-63(uf198)* animals. Points indicate measurements from independent animals. Mean ± SEM are indicated. \*\*\*\**p*<0.0001, student’s t-test. (C)N=12,13. (D)N=11,12. **E,F.** Quantification of the percentage of animals paralyzed following 2 hr exposure to aldicarb (1 mM) (E) or levamisole (250 µM) (F). *unc-63(uf198)* animals are resistant to aldicarb and levamisole. Muscle expression (P*myo-3*::*unc-63 cDNA*) fails to fully restore levamisole sensitivity of *unc-63(uf198)* mutants but is sufficient to restore aldicarb sensitivity. Points indicate independent measurements from trials of at least 10 animals each. Mean ± SEM are indicated. \*\*\*\**p*<0.0001, one-way ANOVA with Tukey’s multiple comparisons test. (E) WT n=4, *acr-12* n=4, *unc-63(uf198)* n=4, *unc-63(uf198);Pmyo-3::unc-63(+)* n=3, *unc-63(x37)* n= 4. (F) WT n=6, *unc-49* n=5, *unc-63(uf198)* n=5, *unc-63(uf198);Pmyo-3::unc-63(+)* n=4, *unc-63(x37)* n= 5, *unc-63(x37);Pmyo-3::unc-63(+)* n=4. **G,H.** Percentage of animals paralyzed over time in the presence of aldicarb (1 mM) (G) or levamisole (250 µM) (H). Heterozygous *unc-63(uf198/+)* animals (grey) treated with either aldicarb or levamisole paralyze more slowly than drug-treated wild type controls (black). Each data point indicates the mean of 4 (G) or 5 (H) independent trials of at least 10 animals each. \**p*<0.05, \*\**p*<0.01, \*\*\**p*<0.001, \*\*\*\**p*<0.0001, two-way ANOVA with Tukey’s multiple comparisons test.

We next sought to investigate the assembly and clustering of L-AChRs at neuromuscular synapses in *unc-63(uf198)* animals. We quantified the clustering of L-AChRs in muscles using muscle-specific expression of UNC-29::GFP, an obligate non-alpha subunit of L-AChRs [21,25]. The number and fluorescence intensity of muscle L-AChR (UNC-29::GFP) clusters were strikingly decreased in *unc-63(uf198)* mutants compared with control (**Fig. 3B-D**). This result suggests that the assembly of muscle L-AChR and GABA neuron ACR-12 AChR complexes are similarly disrupted in *unc-63(uf198)* mutants. However, our observation that movement is not appreciably affected in these animals supports the idea that a low level of receptor assembly persists in *unc-63(uf198)* animals and is sufficient to support muscle contraction and movement.

To further investigate the function of muscle L-AChR in *unc-63(uf198*) animals, we measured the effects of exposure to either the cholinesterase inhibitor aldicarb or the L-AChR agonist levamisole. Wild-type animals are paralyzed in response to prolonged treatment with either chemical. The time course over which paralysis occurs is widely used as a measure of cholinergic function. We found that *unc-63(uf198)* animals are strongly resistant to the paralyzing effects of either aldicarb or levamisole, similar to animals carrying the severe *unc-63(x37)* loss-of-function allele (**Fig. 3E,F**). Muscle-specific expression of wild-type *unc-63* in *unc-63(uf198)* animals restored paralysis in response to aldicarb treatment; however, only weakly altered paralysis in response to levamisole. Expression of the same transgene in *unc-63(x37)* mutants was sufficient to restore levamisole-induced paralysis (**Fig. 3F**). Together, our analysis suggests that the P44L substitution in *unc-63(uf198)* mutants may have dominant negative effects, particularly for activation by levamisole. Consistent with this interpretation, we found that *unc-63(uf198)* heterozygous animals had increased resistance to paralysis by aldicarb and levamisole compared with control (**Fig. 3G,H**), suggesting *unc-63(uf198)* is a semi-dominant allele.

To investigate muscle responses to cholinergic activation in *unc-63(uf198)* mutants, we next measured muscle calcium transients evoked by photostimulation of cholinergic motor neurons. To measure muscle calcium responses, we expressed the genetically encoded calcium sensor GCaMP6s in muscles together with the red-shifted channelrhodopsin Chrimson in cholinergic motor neurons [5]. As the cholinergic synaptic response is mediated through both UNC-63-containing L-AChRs and homomeric ACR-16 AChRs [26,30,31], we pursued these experiments in animals lacking *acr-16*, allowing us to focus on L-AChR-mediated muscle calcium responses. Depolarization of cholinergic motor neurons elicited robust calcium responses that were closely synchronized to the onset of the light stimulus in the muscles of wild-type control animals (response within 200 ms of stimulus onset) (**Fig. 4A,B**). The amplitude of these synchronous calcium responses was dramatically reduced in *unc-63(uf198);acr-16(ok789)* double mutants compared to either wild type or *acr-16(ok789)* single mutants (**Fig. 4A,B**). We observed a similar reduction in these synchronous calcium responses in animals that carried the presumptive null allele *unc-63(x37)* in combination with the *acr-16(ok789)* deletion (**Fig. 4B**). Together, these results suggest that *unc-63(uf198)* is a severe reduction-of-function allele. However, we noted another feature of muscle calcium responses in *unc-63(uf198);acr-16(ok789)* animals, many muscles in *unc-63(uf198);acr-16(ok789)* animals displayed large, delayed calcium transients that were not closely synchronized to the light stimulus (**Fig. 4A, bottom**). These asynchronous responses occurred more frequently in *unc-63(uf198);acr-16(ok789)* muscles compared with either control or *unc-63(x37);acr-16(ok789)* muscles (**Fig. 4C,D**), suggesting this is a property conferred by the UNC-63 P44L substitution.

**Figure 4.**
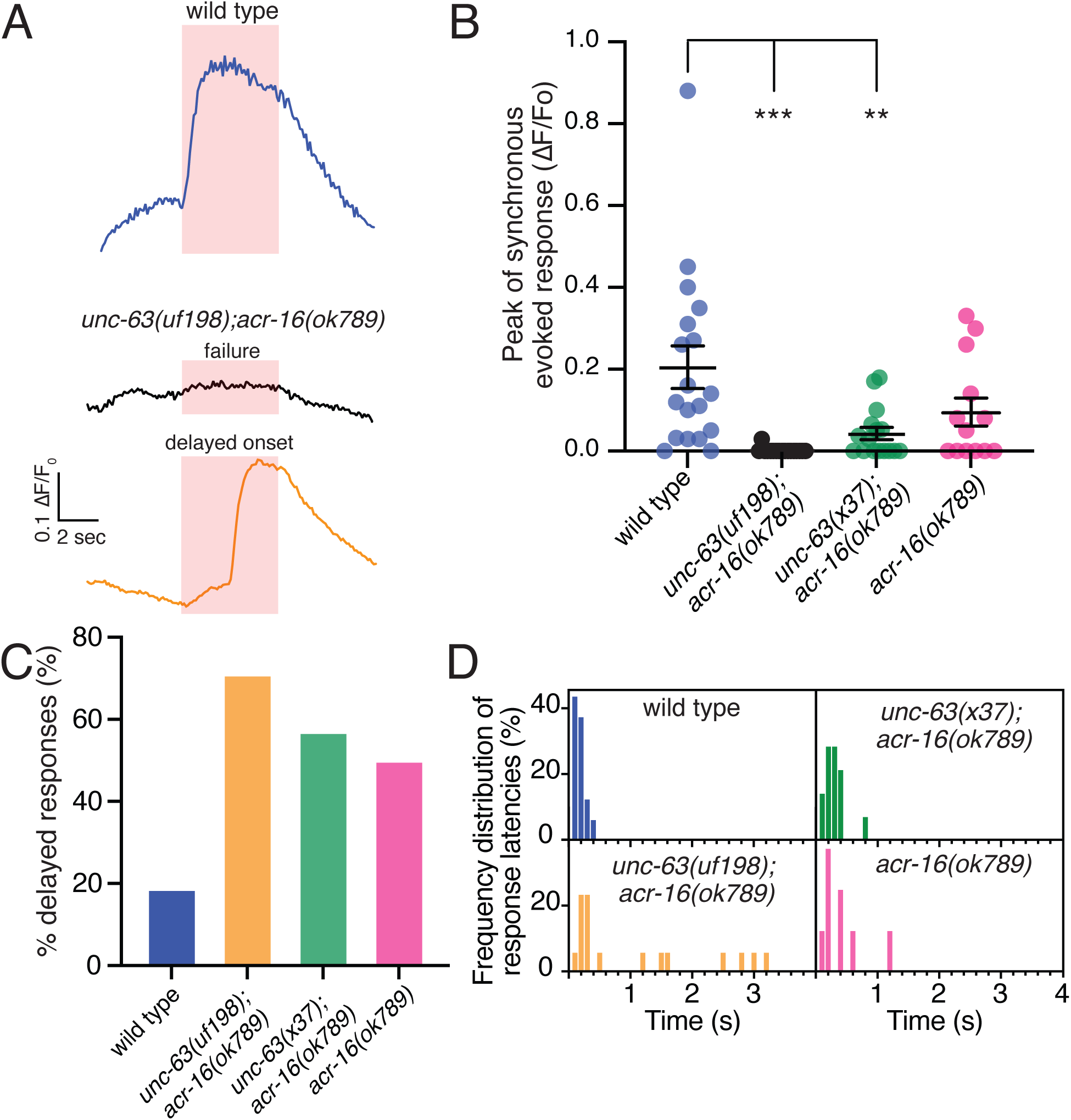
*unc-63(uf198)* muscle responses to cholinergic stimulation are severely disrupted. **A.** Representative evoked muscle calcium responses for wild type and *unc-63(uf198);acr-16(ok789)* animals. Example traces for the two major response classes observed in the *unc-63(uf198);acr-16(ok789)* animals (response failures and delayed onset responses) are shown. Red shading indicates the period of cholinergic photostimulation. **B.** Quantification (mean ± SEM) of the peak muscle responses synchronized to cholinergic depolarization for the indicated genotypes. Points indicate measurements from independent animals. Mean ± SEM are indicated. Responses were scored as synchronous if occurring within 200 ms of stimulus onset. \*\*\**p*<0.001, \*\**p*<0.01, one-way ANOVA with Dunnett’s multiple comparisons test. n=18,17,16,13. **C.** Proportion of delayed onset responses calculated for the indicated genotypes. Note the high proportion of delayed onset responses in *unc-63(uf198)* animals. Responses were classified as delayed if occurring more than 200 ms from the onset of stimulation. Response failures were excluded from the analysis. n=16,16,14,8. **D.** Frequency distribution of response latencies (time from the onset of stimulation to response) for the indicated genotypes for all measurements (including failures). Nearly 90% of wild type muscle calcium responses occurred within 200 ms of stimulus onset. Note the increased response response latencies in *unc-63(uf198)* animals. Response failures were excluded from the analysis. n=16,16,14,8.

### Mutation of *syd-2*/Liprin-**α** disrupts dendritic spine formation

Mapping and sequence analysis of *uf174*, a third mutant isolated from our screen with reduced AChR clusters in GABAergic dendrites, revealed a novel point mutation in the *syd-2* gene (**Fig. 5A**)*. syd-2* encodes the sole *C. elegans* Liprin-α. *syd-2*/Liprin-α was identified as a ligand for LAR family receptor protein tyrosine phosphatases (LAR-RPTP) [32] and implicated as a structural component of active zones based on a series of elegant genetic studies in *C. elegans* [1,10,11]. The SYD-2/Liprin-α family proteins contain a N-terminal coiled-coil region followed by three C-terminal SAM (sterile-α-motif) domains (**Fig. 5A**). The coiled-coil domains are implicated in homo- and hetero-multimerization while the SAM domains form the LH (Liprin Homology) region that binds LAR-RPTP [32]. The *syd-2(uf174)* mutants isolated from the screen carry a C/T missense mutation in exon 18 that produces an early stop (Q1091stop) and truncation of the predicted SAM domain (**Fig. 5A**).

**Figure 5.**
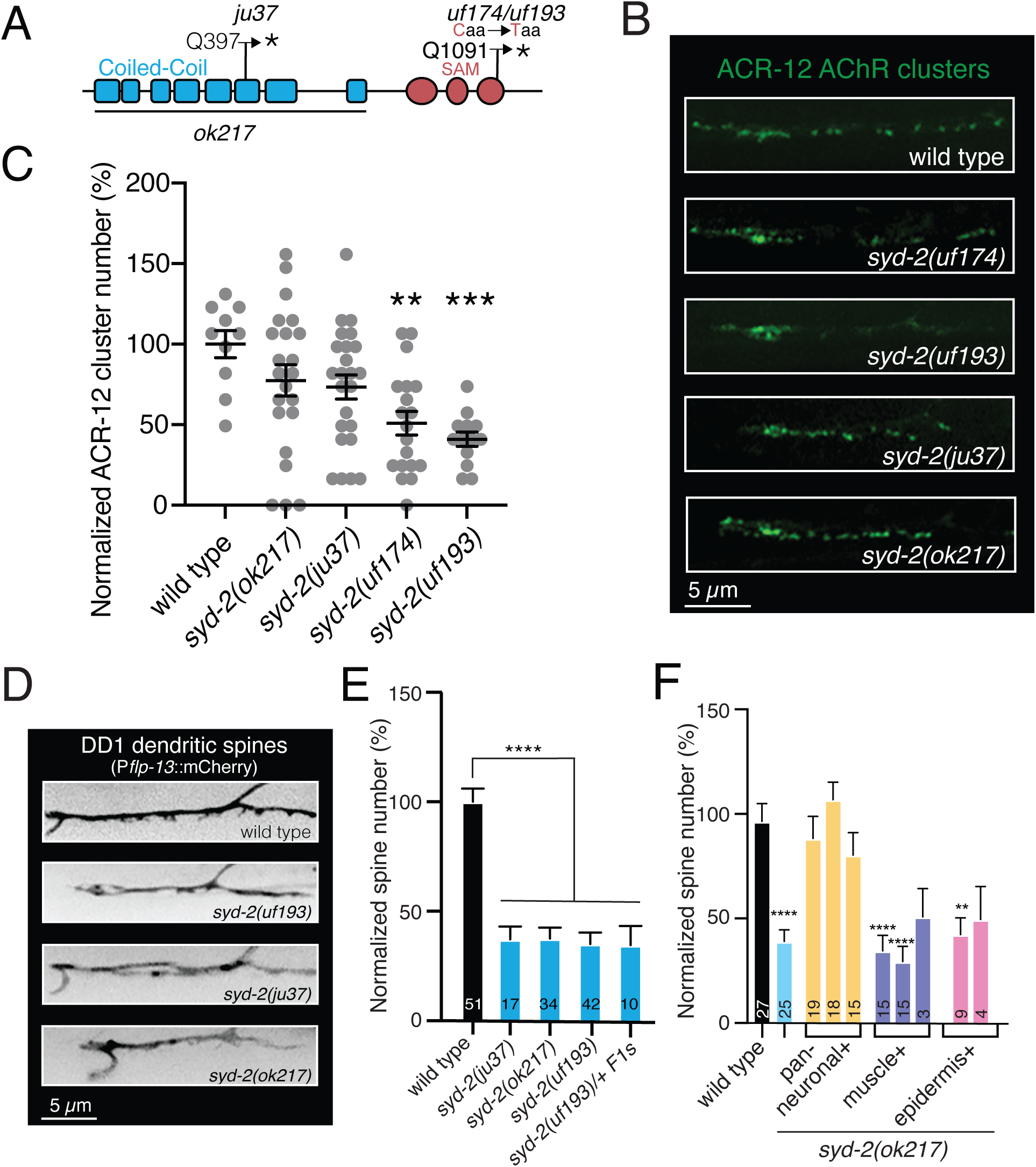
Mutation of *syd-2* disrupts GABAergic dendritic spines. **A.** Domain structure of SYD-2. Coiled-coil (blue) and SAM domains (red) are indicated. Deleted region in *syd-2(ok217)* and approximate positions of mutations in *syd-2(ju37)*, screen isolate *syd-2(uf174)* and CRISPR-engineered *syd-2(uf193)* are indicated. **B.** Confocal images of DD1 ACR-12 AChR localization (P*flp-13*::ACR-12:GFP) in wildtype, screen isolate *syd-2(uf174)*, CRISPR engineered *syd-2(uf193)*, *syd-2(ju37)* and *syd-2(ok217)* animals. **C.** Quantification (mean ± SEM) of ACR-12 AChR clusters in DD GABAergic dendrites of the indicated *syd-2* alleles. \*\**p*<0.01, \*\*\**p*<0.001, one-way ANOVA with Dunnett’s multiple comparison test. N=10,22,24,19,13. **D.** Confocal images of DD1 dendritic spines (P*flp-13*::mCherry) in wild type, CRISPR-engineered *syd-2(uf193),* s*yd-2(ju37)* and *syd-2(ok217)* animals. **E.** Quantification (mean + SEM) of DD1 GABA neuron spine number for the indicated genotypes normalized to wild type. \*\*\*\**p*<0.0001, one-way ANOVA with Dunnett’s multiple comparison test. Numbers in bars indicate the number of animals quantified. **F.** Quantification (mean ± SEM) of GABAergic DD1 dendritic spine number for the genotypes indicated normalized to wild type. Pan-neuronal, but not muscle or epidermal, expression of wild type *syd-2* in *syd-2(ok217)* mutants is sufficient to restore spine number. \*\*\*\**p*<0.0001, \*\**p*<0.01 compared to wild type, one-way ANOVA with Dunnett’s multiple comparison test. Bars indicate independent rescue lines. Numbers in bars indicate number of animals quantified.

ACR-12 AChR clustering was decreased by roughly 50% in *syd-2(uf174)* mutants compared to wild type (**Fig. 5B,C**). To gain support that the mutation in *syd-2* was causal for the disruption in AChR clustering, we engineered the same mutation in otherwise wild-type animals by CRISPR/Cas9 genome editing. CRISPR/Cas9-engineered *syd-2(uf193)* animals showed similar deficits in AChR clustering to those observed in DD neurons of *syd-2(uf174)* isolated from the screen (**Fig. 5B,C**). We next analyzed GABAergic DD dendritic spines (P*flp-13*::mCherry) in *syd-2(uf193)* L4 animals. The number of DD1 dendritic spines in *syd-2(uf193)* animals is decreased by ∼65% compared to wild type animals (**Fig. 5D,E**). Dendritic spine number is also significantly decreased compared to wild type in available *syd-2* loss-of-function mutants, *syd-2(ju37)* and *syd-2(ok217)* (**Fig. 5D,E**). *syd-2(ju37)* animals carry an early stop mutation in sequence encoding the SYD-2 coiled-coil domain while *syd-2(ok217)* animals carry a 2 kb deletion that produces a premature stop [1]. Pan-neuronal expression of wild type *syd-2* cDNA in *syd-2(ok217)* mutants restored spines, indicating a neuronal requirement (**Fig. 5F**). These results suggest that, similar to the synaptic adhesion protein NRX-1 [4,5], SYD-2 may be important for the formation or stabilization of both dendritic spines and ACR-12 AChR clusters at postsynaptic sites. To further investigate the impact of the predicted SYD-2 truncation, we analyzed the number of GABAergic DD dendritic spines in *syd-2(uf193/+)* heterozygotes. Animals heterozygous for *syd-2(uf193)* had decreased spines compared to wild type, similar to homozygous *syd-2(uf193)* mutants and other homozygous loss-of-function alleles tested (**Fig. 5E**), suggesting *uf193* is likely to represent a dominant allele of *syd-2*. SYD-2 has been previously implicated in both presynaptic structural organization and cargo trafficking [1,11,33]. Our findings newly implicate SYD-2 in the formation or stabilization of postsynaptic receptor clusters and dendritic spines, suggesting presynaptic SYD-2 may act indirectly to coordinate postsynaptic organization. Alternatively, it is possible that a postsynaptic SYD-2 pool contributes to spine development and the organization of postsynaptic sites. Additional studies will be required to distinguish between these possibilities.

## Discussion

We pursued an unbiased, genome-wide strategy to elucidate mechanisms directing structural organization of dendrites and postsynaptic specializations localized to dendritic spines in *C. elegans* GABAergic neurons. We identified 3 genes that, when mutated, disrupt the organization of postsynaptic receptor clusters on GABAergic dendrites. These genes define distinct functional pathways required for establishing synaptic connections: (1) the RUN domain cargo adaptor UNC-14 is required for dendrite outgrowth, (2) the nAChR subunit UNC-63 is an obligate subunit of postsynaptic receptors and required for receptor assembly, and (3) the synaptic scaffold SYD-2 is required for spine outgrowth or maintenance.

The *unc-14* allele we obtained from our screen generates an early stop codon and is a predicted null. Prior studies of *unc-14* null mutants demonstrated an interaction between UNC-14 and the the S/T kinase UNC-51 and implicated UNC-14 in axonal elongation as well as axonal delivery of a variety of cargoes including synaptic vesicles, the axon guidance cue UNC-6/Netrin, and the Netrin receptor UNC-5 [6-8,18]. Our analysis of dendritic structure revealed striking morphological abnormalities in GABAergic DD dendrites in addition to the disruption of ACR-12/AChR clustering identified from our initial screen. We are therefore unable to distinguish whether the defects in postsynaptic organization we observed reflect a requirement for UNC-14 in trafficking post-synaptic components, arise secondarily to a failure in the extension and fasciculation of dendritic processes within the ventral nerve cord, or a combination of both. Similar to previous findings for *unc-14(e57)* mutants, we observed bulbous regions in the dendrites of DD GABAergic neurons in *unc-14(uf198)* animals that may represent foci where dendritic material, including post-synaptic proteins, has accumulated due to a failure in trafficking. However, *unc-14* is known to be broadly expressed in neurons [6]. It is therefore also possible that some of the phenotypes we observed in GABAergic DD dendrites reflect a parallel requirement for *unc-14* in the organization of presynaptic cholinergic axons. Consistent with this hypothesis, we found that *unc-14* expression using either an ∼2.9 kb *unc-14* promoter fragment or a pan-neuronal promoter was sufficient to rescue the uncoordinated movement of *unc-14* mutants. In contrast, specific expression in either cholinergic or GABAergic motor neurons was not sufficient to rescue, most consistent with a requirement for *unc-14* expression in both neuron classes for proper motor function.

The P/L substitution mutation in *unc-63* that we isolated disrupts cholinergic receptor clustering in both GABAergic dendrites (P*flp-13*::ACR-12::GFP) and muscles (P*myo-3*::UNC-29::GFP). The mutation, however, does not affect the formation or maintenance of GABAergic dendritic spines, consistent with prior work demonstrating spine outgrowth proceeds independently of receptor clustering [4,5]. Prior structural studies of acetylcholine binding proteins from the freshwater snail, *Lymnaea stagnalis* (*L*-AChBP), saltwater mollusk *Aplysia californica*, and the extracellular ligand binding domain of the mouse muscle alpha1 subunit place this residue adjacent to the N-terminal α-helical region at the apical surface of the receptor [34-36]. Prior mutagenesis studies of chimeric mouse AChRs have implicated this region in receptor assembly and ligand sensitivity [29], consistent with our findings for the P/L substitution in UNC-63. In addition to available null alleles, *C. elegans* mutations that alter UNC-63 subunit functional properties have been characterized previously. For instance, *unc-63(x26)* (C151Y) disrupts the Cys-loop region, a motif important for forming the agonist binding site, and causes a decrease in channel opening frequency [37]. However, *unc-63(x26)* is thought to exclusively alter channel function, not expression. Our analysis suggests that the conserved proline which is altered in *unc-63(uf174)* and *unc-63(uf198)* animals is important for proper assembly and function of UNC-63-containing AChRs in both muscles and GABAergic dendrites. Although the P44L substitution is not at the ligand binding Cys-loop motif, the proline to a leucine change may alter the structure of the extracellular ligand binding domain, impacting the both assembly/folding of UNC-63 subunit and ligand binding.

Given the severe disruption in nAChR clustering we observed and the resistance to pharmacological activation by aldicarb or levamisole, it is surprising that the movement of *unc-63(uf198)* animals is not more severely disrupted. Our studies of evoked calcium responses in muscles indicate that calcium responses do sometimes occur in muscles of *unc-63(uf198)* animals, but these responses are not strictly time-locked to the stimulus. The most parsimonious interpretation of our findings is that the allele is a strong reduction-of-function, but allows sufficient residual receptor activity to support movement. One possibility is that inefficient activation of the receptor may over time produce muscle depolarization. Alternatively, the mutation may indirectly lead to increased spontaneous muscle activity, though a mechanism to account for such an effect remains unclear. Mammalian ionotropic AChRs are linked with myasthenic syndromes and nicotine addiction [38] while nematode AChRs are common targets for anthelmintic compounds [39]. Our identification and characterization of *unc-63(uf198)* animals therefore offer additional tools for comparative structure-function studies of this conserved receptor family.

The isolation of a novel *syd-2* allele from our screen newly highlights the importance of this scaffold protein for maintaining postsynaptic organization. Evidence from prior studies across a variety of models implicates SYD-2/Liprin-α as one of several core scaffold proteins responsible for orchestrating presynaptic active zone organization. SYD-2/Liprin-α and the RhoGAP domain and PDZ domain-containing scaffold protein, SYD-1, are suggested to localize to nascent active zones in the early stages of presynaptic development, and to subsequently recruit additional presynaptic proteins for presynaptic assembly [11,40-48]. In particular, SYD-1 is suggested to function upstream of SYD-2 and to promote SYD-2 localization and activity at presynaptic sites [11,49]. Interestingly, *Drosophila dSyd1* and *neurexin (dnrx)* mutants have similar active zone defects and form an *in vivo* complex *where* dSyd1 stabilizes neurexin at active zones through PDZ domain interactions [11,49,50]. Similarly, in mammalian neurons, Liprin-α forms a complex with Neurexin1 and another presynaptic scaffold protein Mint1 to regulate the stability of Neurexin1 [51]. We showed previously that presynaptic NRX-1/neurexin is required for the maintenance of developing spines [4,5], raising the possibility that the postsynaptic alterations we observe in *syd-2* mutants arise secondarily to a mislocalization of NRX-1 at presynaptic sites. Alternatively, our findings may indicate a postsynaptic requirement for SYD-2. Though less well-characterized compared with its presynaptic roles, immuno-EM studies of rodent cortical neurons have demonstrated that Liprin-α is also localized to postsynaptic sites [52]. Postsynaptic Liprin-α has been implicated in AMPAR clustering and in shaping morphological features of dendritic spines through interactions with GRIP1 (glutamate receptor-interacting protein 1) and LAR family of receptor protein tyrosine phosphatases [52-54]. Similarly, postsynaptic Liprin-α organizes AChRs at the mouse neuromuscular junction [55]. In *C. elegans*, current evidence supports predominant presynaptic SYD-2 localization [1,11]. Nonetheless, additional studies employing strategies for cell-specific inactivation or deletion of *syd-2* will be required to cleanly dissect potential pre-versus postsynaptic roles at synapses onto GABAergic neurons.

Regardless of the precise mechanism of action, the *syd-2(uf174)*/*syd-2(uf193)* alleles offer additional tools for functional analysis of SYD-2 at synapses. The mutation we isolated generates an early stop codon in the region encoding the predicted SYD-2 SAM domain and has similar impacts on spines in *syd-2(uf193)* homozygotes and *syd-2(uf193/+)* heterozygotes, indicating the mutation has dominant effects. Prior work identified a putative *syd-2* gain-of-function mutation, *syd-2(ju487)*, that also has dominant effects. *syd-2(ju487)* carries a missense mutation that produces an R/C substitution at position 184 (R184C) of the conserved Liprin Homology 1 (LH1) domain [11]. This mutant has synaptic phenotypes opposite to those associated with *syd-2* loss-of-function alleles, including elongated dense core projections and hypersensitivity to the acetylcholine esterase inhibitor aldicarb [11,56]. In contrast, our analysis of *syd-2(uf193/+)* demonstrates this dominant mutation phenocopies known loss-of-function mutants. Our findings suggest the possibility that a dominant-negative, truncated SYD-2 protein is produced in which the SAM domain is disrupted, perhaps locking SYD-2 in an inactive conformation or interfering with interactions with activating proteins [10,57]. Our observation that the reduction in ACR-12 AChR clusters in the *syd-2* mutant isolated from our screen is more severe compared with other available loss-of-function alleles may also point toward a dominant negative role.

Collectively, our studies of *unc-14, unc-63,* and *syd-2* offer new avenues for understanding synapse development and organization. Moreover, the novel alleles we isolated from our screen provide new tools for gaining additional insights into the functions of these conserved proteins at synapses.

## Materials and Methods

### Strain maintenance

All strains used are derivatives of N2 Bristol and maintained at room temperature. Animals were grown on nematode growth media plates (NGM) seeded with *E. coli* strain OP50. A complete list of all strains used are found in Table 1.

### EMS mutagenesis, whole-genome sequencing and bioinformatic data analysis

To identify genes important for the development or maintenance of synapses onto GABAergic neurons, we conducted a semi-clonal F2 forward screen [58,59] to obtain mutants where the clustering of GFP-tagged ACR-12 receptors is disrupted. Transgenic animals stably expressing P*flp-13*::ACR-12::GFP [*ufIs126*] were mutagenized for 4 hours using 25 µM ethyl methanesulfonate (EMS) following standard protocols [60]. Mutagenized L4/young adults were transferred to NGM plates. 3-4 days later, F1s were transferred (2/plate) to 50 plates (100 F1s). From each of the 50 F1 plates, 8 F2s were transferred to 4 new plates (2/plate) yielding 400 F2s on 200 plates. L4 stage F2 progeny were screened for disruptions in ACR-12::GFP clustering in DD neurons (absent, displaced, ectopic, decrease or increased number) by spinning disk confocal microscopy. Roughly 3300 haploid genomes were screened in total.

Candidate mutants were mapped by variant discovery mapping as previously described [61,62]. Briefly, mutant phenotypes were first confirmed in subsequent generations. Confirmed candidates were then backcrossed to the starting, non-mutagenized strain carrying P*flp-13*::ACR-12::GFP together with a visible marker used in the crosses. Homozygous F2 progeny from 16-23 backcrossed recombinants were pooled for DNA extraction and whole genome sequencing (Novogene). Homozygous variants within the mapped region were included in the final candidate list. Raw sequence data was processed using Galaxy/Cloudmap mapping, subtracting, and filtering pipelines [63]. Sequence reads were aligned to the reference genome (wild type/N2 Bristol WS220) [64] using the fast-short read Burrows-Wheeler Aligner (BWA). For high coverage and high-quality datasets only reads with a Phred score of >30 (according to FastQC) for each base pair was used. Variants were identified using GATTK Unified Genotyper from the GATK suite. Variations present in the P*flp-13*::ACR-12::GFP starting strain were subtracted and filtered (read depth of ≥3) from the mutant sequence results using GATK SelectVariants from the GATK suite. Variants were plotted using CloudMap and annotated using SnpEff.

### Imaging and fluorescence quantification

L4 or Day 1 adult animals were immobilized with sodium azide (0.3 M) on 2% or 5% agarose pads. Z-stack confocal volumes were obtained using a Yokogawa CSU-X1 spinning disk confocal mounted on an Olympus BX51WI upright microscope equipped with a Hamamatsu EM-CCD camera and 63x oil immersion objective. Analysis of synapse number and fluorescence intensity was performed using FIJI ImageJ software [65] using defined threshold values acquired from control experiments for each fluorescent marker. Statistical comparisons for all synaptic and spine analyses were performed by student’s t-test for experiments with two groups, or by one-way or two-way ANOVA with the appropriate post-hoc test for experiments with multiple groups.

### Molecular biology

Plasmids were constructed using the two-slot Gateway Cloning system (Invitrogen) and confirmed by restriction digest and sequencing. All plasmids and primers used in this work are described in Table 2.

### CRISPR/Cas9 design and gene editing

The Bristol N2 strain was used as the wild-type strain for all CRISPR/Cas9 editing. *unc-63* and *syd-2* gene editing was performed by CRISPR/Cas9 using established protocols [66]. Site-specific sgRNAs and repair template donor ssODNs were manually designed and generated by IDT (Integrated DNA Technologies, Inc.). Mutations in the *dpy-10* gene were used as a CRISPR co-conversion marker. Base substitutions were confirmed via Sanger sequencing. All crRNAs, ssODNs, and PCR screening sequences are reported in Table 2.

### Aldicarb and levamisole assays

Assays were performed on synchronous populations of Day 1 adult animals. At least 10 worms of each genotype were transferred to separate plates containing NGM agar and either 1 mM aldicarb or 250 μM levamisole. The number of paralyzed animals was recorded every 15 minutes and the rate of paralysis was plotted as a function of time using GraphPad Prism. Assays were repeated on at least 3 separate days.

### Behavioral assays

Staged Day 1 adult animals were placed on behavioral assay plates coated with a thin OP50 lawn. Following one minute of habituation, worm movement was recorded using the WormLab Imaging System (MBF Bioscience). Post-hoc analysis was performed using WormLab software. The total length of the worm’s movement path (track length), calculated by summing the lengths of all forward and backward movements, over 1-minute duration from the start of image acquisition was calculated. Statistical comparisons were performed by one-way ANOVA with Bartlett’s correction using GraphPad Prism.

### Calcium imaging

Simultaneous optogenetics and calcium imaging was performed as previously described [5]. Ca^2+^ transients were recorded from muscle cells in transgenic animals co-expressing P*myo-3*::NLSwCherry::SL2::GCaMP6s (from M. Alkema) along with P*acr-2*::Chrimson. Animals were placed on plates seeded with OP50 containing 2.75 mM All-Trans Retinal (ATR) for 24 hr prior to experiments. Young adults were immobilized on 5% agarose pads in 2,3-Butanedione monoxime (BDM) (30 mg/mL). For all genotypes, control animals grown in the absence of ATR were imaged. Data were acquired at 10 Hz sampling using Volocity software. Pre-stimulus baseline fluorescence (F0) was calculated as the average of the corrected background-subtracted data points in the first 4 s of the recording (prior to stimulus onset). The corrected fluorescence response data was normalized to prestimulus baseline as ΔF/F0, where ΔFL=LF– F0. Peak ΔF/F0Lwas determined by fitting a Gaussian function to the ΔF/F0 Ltime sequence using Multi peak 2.0 (Igor Pro, WaveMetrics). All data collected were analyzed, including failures (no response to stimulation). All responses occurring within the stimulation window (5 s) that could be fitted with a Gaussian function were scored as positive and quantified. LLatency was calculated as the time after stimulus onset for the fluorescence intensity to reach two times the pre-stimulus baseline standard deviation [67].

## Supplementary Material and Tables

**Table 1.** Strains used in this study.

| GENOTYPE | STRAIN | DESCRIPTION |
| --- | --- | --- |
| *ufls126* | IZ1905 | *Pflp-13::ACR-12::GFP* (pCL32 @ 150 ng/μL) + *Plgc-11::mCherry* @ 20 ng/μL |
| *ufls128* | IZ1464 | *Pflp-13::mCherry* (pAP31 @ 50 ng/μL) + *Plgc-11::mCherry* @ 20 ng/μL |
| *unc-14(uf169)* 10X outcross | IZ3020 | Outcrossed to N2 Bristol |
| *unc-14(uf169); ufls126* | IZ2498 |  |
| *unc-14(uf169); ufls128* | IZ2802 |  |
| *unc-14(uf169); ufls126; ufEx1069* | IZ2790 | *ufEx1069* = PCR amplified genomic *unc-14* rescue + *Punc122::GFP* @ 50 ng/μL |
| *unc-14(uf169); ufEx1922* | IZ4534 | *ufEx1922* = *Prgef-1::unc-14* cDNA (pJR14 @ 10 ng/μL + *Pinx-6::GFP* (pDO125 @ 50 ng/μL) |
| *unc-14(uf169); ufEx1156* | IZ2932 | *ufEx1156* = *Punc-47::unc-14* cDNA-*yk1203h04* (pDO17 @ 10 ng/μL) + *Punc-122::GFP* @ 50 ng/μL |
| *unc-14(uf169); ufEx1173* | IZ2963 | *ufEx1173* = *Punc-17Beta::unc-14* cDNA (pDO15 @ 10 ng/μL) + *Punc-122::GFP* @ 50 ng/μL |
| *unc-14(e57)* | CB57 |  |
| *unc-63(uf175); ufls126* | IZ2438 | Screen isolated mutant allele |
| *unc-63(uf198)* | IZ3558 | CRISPR generated mutant allele |
| *unc-63(x37)* | IZ231 |  |
| *unc-63(ok1075)* | VC731 | CGC |
| *acr-16(ok789)* | IZ57 |  |
| *unc-63(uf198); acr-16(ok789)* | IZ4505 |  |
| *unc-63(x37); acr-16(ok789)* | IZ60 |  |
| *acr-12(ok367)* | IZ241 |  |
| *unc-49(e382)* | CB382 | CGC |
| *unc-63(uf198); ufls126* | IZ3917 |  |
| *unc-63(ok1075); ufls126* | IZ1427 |  |
| *unc-63(uf175); ufls126; ufEx1104* | IZ2845 | *ufEx1104* = *Punc-47::unc-63* cDNA (pAP59 @ 30 ng/μL) + *Punc-122::GFP* @ 50 ng/μL |
| *unc-63(uf175); ufls126; ufEx1105* | IZ2848 | *ufEx1105* = *Punc-47::unc-63* cDNA (pAP59 @ 30 ng/μL) + *Punc-122::GFP* @ 50 ng/μL |
| *unc-63(uf175); ufls126; ufEx1111* | IZ2855 | *ufEx1111* = *Punc-47::unc-63* cDNA (pAP59 @ 30 ng/μL) + *Punc-122::GFP* @ 50 ng/μL |
| *unc-63(uf198); ufls128* | IZ3714 |  |
| *unc-63(uf198); ufls2* | IZ3753 | *ufls2* = *Pmyo-3::unc-29::GFP* (pDM956) + *lin-15(+)* (pJM23) |
| *unc-63(uf198); ufEx1907* | IZ4494 | *ufEx1907* = *Pmyo-3::unc-63* cDNA (pAP212 @ 25 ng/μL) + pDO125 @ 50 ng/μL |
| *unc-63(x37); ufEx1907* | IZ4614 | *ufEx1907* = *Pmyo-3::unc-63* cDNA (pAP212 @ 25 ng/μL) + pDO125 @ 50 ng/μL |
| *zfEx813; ufls157* | IZ4568 | *zfEx813* (Mark Alkema) = *Pmyo-3::NLSwCherry::SL2::GCaMP6s* (50 ng/μL); *ufls157* = *Pacr-2::Chrimson* (pRB2 @ 50 ng/μL) |
| *unc-63(uf198); acr-16(ok789); zfEx813; ufls157* | IZ4607 |  |
| *unc-63(x37); acr-16(ok789); zfEx813; ufls157* | IZ4596 |  |
| *acr-16(ok789); zfEx813; ufls157* | IZ4611 |  |
| *syd-2(uf174)* | IZ3341 | Screen isolated mutant allele |
| *syd-2(uf193)* | IZ3356 | CRISPR generated mutant allele |
| *syd-2(ok217)* | ZM607 | CGC |
| *syd-2(ju37)* | CZ900 | CGC |
| *syd-2(uf174); ufls126* | IZ2321 |  |
| *syd-2(uf193); ufls126* | IZ3419 |  |
| *syd-2(ju37); ufls126* | IZ3028 |  |
| *syd-2(ok217); ufls126* | IZ2984 |  |
| *syd-2(uf193); ufls128* | IZ3429 |  |
| *syd-2(ju37); ufls128* | IZ3043 |  |
| *syd-2(ok217); ufls128* | IZ2983 |  |
| *syd-2(ok217); ufls128; ufEx1430* | IZ3424 | *ufEx1430* = *Prgef-1::syd-2 cDNA* (pDO63 @ 10 ng/μL) + *Punc-122::GFP* @ 50 ng/μL |
| *syd-2(ok217); ufls128; ufEx1342* | IZ3266 | *Pmyo-3::syd-2cDNA* (pDO33 @ 10ng/μL) + *Punc-122::GFP* @ 50 ng/μL |
| *syd-2(ok217); ufls128; ufEx1343* | IZ3267 | *Pelt-3::syd-2cDNA* (pDO36 @ 10ng/μL) + *Punc-122::GFP* @ 50 ng/μL |

**Table 2.** Plasmids used in this study.

|  |  |  |
| --- | --- | --- |
| *Pflp-13::ACR-12::GFP* | pCL32 | Linear recombination by Gateway of pENTR-3'-*Pflp-13* and pDest-38 to generate *Pflp-13::ACR-12::GFP* |
| *Pflp-13::mCherry* | pAP31 | Linear recombination by Gateway of pENTR-5'-*Pflp-13* and pDest-17 to generate *Pflp-13::mCherry* |
| *unc-14* promoter | pENTR-5'-22 | For *unc-14* rescue, the *unc-14* genomic region including ~2.9 kb upstream from the ATG was amplified from N2 genomic DNA using primers OMF2079 and OMF2080, and ligated into pENTR D-TOPO using CACC overhang ligation to create pENTR-5'-22. |
| Pan-neuronal rescue of *unc-14* | pJR14 | Linear recombination by Gateway of pENTR-3'-*Prgef-1* and pDest-141 to generate *Prgef-1::unc-14 cDNA* |
| GABA rescue of *unc-14* | pDO17 | Linear recombination by Gateway of pENTR-3'-*Punc-47* and pDest-141 to generate *Punc-47::unc-14 cDNA* |
| Cholinergic rescue of *unc-14* | pDO15 | Linear recombination by Gateway of pENTR-3'-*Punc-17β* and pDest-141 to generate *Punc-17β::unc-14 cDNA* |
| *unc-63 cDNA* | pDest-57 | *unc-63* amplified from pTB212 using primers OMF1120 & OMF1121, and ligated into pDest-38 using NheI & SacI |
| GABA rescue of *unc-63* | pAP59 | Linear recombination by Gateway of pENTR-3'-*Punc-47* and pDest-57 to generate *Punc-47::unc-63 cDNA* |
| Muscle rescue of *unc-63* | pAP212 | Linear recombination by Gateway of pENTR-3'-*Pmyo-3* and pDest-57 to generate *Pmyo-3::unc-63 cDNA* |
| Cholinergic Chrimson plasmid | pRB2 | Linear recombination by Gateway of pENTR-5'-*Pacr-2* and pDest-104 to generate *Pacr-2::Chrimson* |
| *syd-2 cDNA* | pDest-155 | *syd-2 cDNA* was RT-PCR amplified from N2 RNA using primers OMF2336 and OMF2337, and ligated into pDest-149 to create pDest-155 |
| *syd-2 cDNA* | pDest-159 | *syd-2 cDNA* was RT-PCR amplified from N2 RNA using primers OMF2336 and OMF2337, and ligated into NheI/NgoMIV digested pDest-150 to create pDest-159 |
| *Prgef-1::syd-2 cDNA* | pDO63 | Linear recombination by Gateway of pENTR-5'-*Prgef-1* and pDest-155 to generate *Prgef-1::syd-2 cDNA* |
| *Pmyo-3::syd-2 cDNA* | pDO33 | Linear recombination by Gateway of pENTR-3'-*Pmyo-3* and pDest-159 to generate *Pmyo-3::syd-2 cDNA* |
| *Pelt-3::syd-2 cDNA* | pDO36 | Linear recombination by Gateway of pENTR-34 and pDest-159 to generate *Pelt-3::syd-2 cDNA* |

**Table 3.**
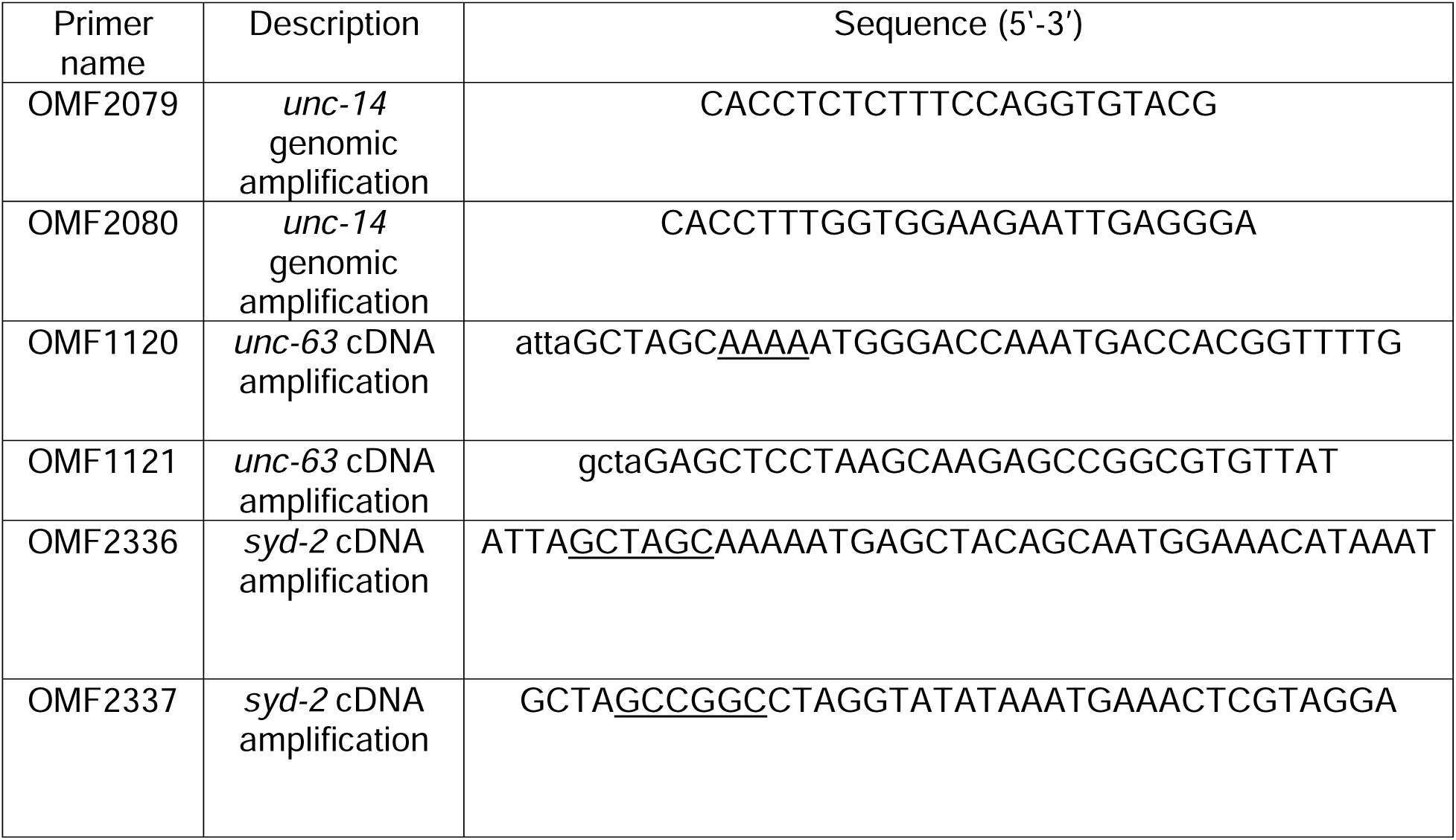
Primers used in this study.

## Supplementary Information on generated CRISPR strains

All CRISPR/Cas9 used in these studies were generated as described [66].

### unc-63(uf198)

Target sequences were selected on exon 2 of *unc-63*. Forward and reverse oligonucleotides were designed to contain the target sequence and overhangs compatible with *Bsa*I sites in plasmid Forward and reverse oligonucleotides were designed to contain the target sequence and overhangs compatible with *BsaI* sites in plasmid pPP13, a modified version of pRB1017 where the sgRNA scaffold was replaced with the sgRNA(F+E) scaffold [68]. Forward and reverse oligonucleotides were annealed and ligated into pPP13 cut with *Bsa*I to create the gRNA plasmids. Plasmids were confirmed by sequencing with M13 reverse primer. A DNA mixture of pDD162 (P*eft-3*::Cas-9) (50 ng/µL), the gRNA plasmids (40 ng/µL each), pPP15 (50 ng/µL) (*dpy-10* target, a modified version of pJA58 were the sgRNA scaffold was replaced with the sgRNA(F+E) scaffold, the ssODN repair template for *dpy10(cn64)* (20 ng/µL) and the ssODN repair template for *unc-63* (Donor uf175) (80 ng/µL) was prepared in injection buffer and injected into N2 worms. Donor *uf175* contains a point mutation cCg/cTg resulting in a P/L amino acid change and a point mutation to eliminate an *AgeI* site for screening of mutants. Mutations in the *dpy-10* gene were used as a CRISPR co-conversion marker. The F1 progeny were screened for Rol and Dpy phenotypes 3-4 days after injection and then for the target mutation in the *unc-63* coding region by PCR and *AgeI* digest. The *unc-63(uf198)* mutant contains a cCg/cTg point mutation at codon 44 P/L.

### syd-2(uf193)

Target sequences were selected on exon 18 of *syd-2*. Forward and reverse oligonucleotides were designed to contain the target sequence and overhangs compatible with *Bsa*I sites in plasmid pPP13, a modified version of pRB1017 where the sgRNA scaffold was replaced with the sgRNA(F+E) scaffold [68]. Forward and reverse oligonucleotides were annealed and ligated into pPP13 cut with *Bsa*I to create the gRNA plasmids. Plasmids were confirmed by sequencing with M13 reverse primer. A DNA mixture of pDD162 (P*eft-3*::Cas-9) (50 ng/µL), the gRNA plasmids (50 ng/µL each), pPP15 (50 ng/µL) (*dpy-10* target, a modified version of pJA58 were the sgRNA scaffold was replaced with the sgRNA(F+E) scaffold, the ssODN repair template for *dpy10(cn64)* (20 ng/µL) and the ssODN repair template for *syd-2* (Donor uf174) (50 ng/µL) was prepared in injection buffer and injected into N2 worms. Donor *uf174* contains a point mutation Caa/Taa resulting in a Q/stop change and the generation if an *AflII* site for screening of mutants. Mutations in the *dpy-10* gene were used as a CRISPR co-conversion marker. The F1 progeny were screened for Rol and Dpy phenotypes 3-4 days after injection and then for the target mutation in the *syd-2* coding region by PCR and *AgeI* digest. The *syd-2(uf193)* mutant contains a point mutation CCA/TAA resulting in a Q/stop at codon 1091.

## Acknowledgments

We thank Christopher Lambert and Julia Russell for help with molecular biology and strain building, and Andrea Thackeray for help with the EMS mutagenesis screen. CRISPR/Cas9 engineered mutations were designed and produced by Paola Perrat and Peng Ding in the University of Massachusetts Chan Medical School CRISPR Core Facility. Some strains were provided by the CGC, which is funded by NIH Office of Research Infrastructure Programs (P40 OD010440). This research was supported by NINDS RO1NS064263 (MMF), NSF IOS1755019 (MMF), NSF IOS1754986 (MLL) and NIH F31NS103365 (DBO).

## Author contributions

DBO directed the screen and mutant isolation, coordinated whole genome sequencing (WGS) efforts, identified the causal mutations, and performed a majority of the functional characterization of the candidates. SR performed Ca^2+^ imaging experiments/analysis and assisted with preparing the manuscript. KB performed strain building, behavior experiments, imaging and analysis. CB helped with screen design, and with MD performed bioinformatics analysis on WGS results. HK and NG built strains, performed behavior experiments and analysis. MLL performed behavior experiments and data analysis. MMF was responsible for project conceptualization, guidance, and writing the manuscript.

